# A dynamic power-law sexual network model of gonorrhoea outbreaks

**DOI:** 10.1101/322875

**Authors:** Lilith K Whittles, Peter J White, Xavier Didelot

## Abstract

Human networks of sexual contacts are dynamic by nature, with partnerships forming and breaking continuously over time. Sexual behaviours are also highly heterogeneous, so that the number of partners reported by individuals over a given period of time is typically distributed as a power-law. Both the dynamism and heterogeneity of sexual partnerships are likely to have an effect in the patterns of spread of sexually transmitted diseases. To represent these two fundamental properties of sexual networks, we developed a stochastic process of dynamic partnership formation and dissolution, which results in power-law numbers of partners over time. Model parameters can be set to produce realistic conditions in terms of the exponent of the power-law distribution, of the number of individuals without relationships and of the average duration of relationships. Using an outbreak of antibiotic resistant gonorrhoea amongst men have sex with men as a case study, we show that our realistic dynamic network exhibits different properties compared to the frequently used static networks or homogeneous mixing models. We also consider an approximation to our dynamic network model in terms of a much simpler branching process. We estimate the parameters of the generation time distribution and offspring distribution which can be used for example in the context of outbreak reconstruction based on genomic data. Finally, we investigate the impact of a range of interventions against gonorrhoea, including increased condom use, more frequent screening and immunisation, concluding that the latter shows great promise to reduce the burden of gonorrhoea, even if the vaccine was only partially effective or applied to only a random subset of the population.

## Introduction

In 2017 the WHO added *Neisseria gonorrhoeae* to its priority list of bacterial pathogens in response to the global spread of antibiotic resistance [1]. The bacteria have developed resistance to every therapy used against them, from penicillin through to third-generation cephalosporins [2]. At a time when resistance to first line therapies is increasingly observed [3], it is more important than ever to understand the transmission dynamics of the infection, and how interventions might be used to reduce the burden on antibiotic treatment [4].

It has been well documented that heterogeneity in sexual activity levels has an impact on disease transmission, with individuals who have many partners bearing much of the burden of disease [5–8]. However, the risk of acquiring and passing on a sexually transmitted infection (STI) depends not only on an individual’s sexual risk profile, but also on their position in the wider sexual network [9,10]. Furthermore the structure of an underlying network affects the probability that an infection that is introduced to the network leads to an outbreak, as well as the size and longevity of any outbreaks that occur [11,12]. As such, it is important to take into account the structure of the underlying sexual network when modelling STI outbreaks.

The distribution of the number of sexual contacts within a network is known as its degree distribution. Several studies have shown that real world sexual networks often have degree distributions that obey a power-law [13–15], where the probability of having *k* partners over a given period of time is proportional to *k*^−γ^. The constant γ is usually between 1 and 4, and different values have been observed in heterosexual and same-sex networks, as well as between genders [14]. Power-law networks exhibit high levels of heterogeneity, with the majority of individuals having a relatively small number of contacts, while a few have many. The standard method to simulate power-law networks is to use a system of preferential attachment, in which individuals are added one by one, connecting with a higher probability with existing individuals who already have a large number of partners [16]. Once all individuals have been added, the network thus created is guaranteed to have a static power-law distribution. It is important to note that even though the preferential attachment algorithm is dynamic in nature, the dynamic method used is purely a technique for generating a static network and does not in any way reflect the dynamics known to occur in real world sexual networks. Furthermore, transmission of infection occurs only once the network has been generated, with all partnerships being in place constantly from the beginning of the simulation of infection transmission until the end.

An alternative method of producing static networks with a power-law degree distribution has been proposed based on each network node having an intrinsic fitness parameter, and a function that determines the probability that a network connection exists between any two nodes depending on their fitness [17,18]. In a sexual network, this can be thought of as each individual having an inherent propensity to seek new partnerships, with the probability of occurrence of each possible partnership depending on the mutual attraction of two individuals.

In a sexual network model, the rate of transmission to an uninfected individual depends on the infection status of their sexual partners, rather than on the prevalence of infection in the pool of potential partners, as in compartmental models. Compartmental models that do not explicitly represent partnerships have been shown to underestimate the importance of core groups of highly sexually active individuals in sustaining STI transmission, while overestimating the contribution of long-term partnerships and low-activity individuals [19]. Furthermore, several studies have shown that, in order to explain observed patterns of infection, it is important to take into account not only the network structure but also the duration of partnerships, and the gaps between them [20,21]. It may therefore be necessary to use a dynamically evolving network to correctly simulate the spread of STI outbreaks. However, no algorithm has yet been proposed to simulate a dynamic sexual network with the correct real-world properties such as a power-law distribution of number of sexual partners, so that the difference between such a realistic dynamic network and a more approximate static network has not been assessed.

Here we present a novel approach to dynamic network simulation using stochastic partnership formation and breakdown based on individuals’ intrinsic properties. We demonstrate that our method produces power-law networks, and that it can simulate a population reflecting the observed network characteristics in UK men who have sex with men (MSM). We then simulate an outbreak of gonorrhoea in three types of network: a fully-connected static network, a heterogeneous static network, and our novel heterogeneous dynamic network, showing important differences between all three models in terms of the resulting patterns of transmission. We estimate the resulting offspring distribution (the number of secondary cases caused by each primary case) and generation time distribution (time between infection of a primary case and infection of a secondary case) to assess the likelihood of super-spreading events predicted by each network structure, as well as the predicted duration of infection. We also compare the impact of the number of sexual partnerships on the probability of infection and transmission under each network structure, providing a basis for risk assessment. Finally, using the dynamic network model we investigate the impact of a range of interventions against gonorrhoea, including increased condom use, more frequent screening and a hypothetical vaccine.

## Results

### Analysis of survey data on number of partners

We first analysed the number of partners reported by MSM in the third National Survey of Sexual Attitudes and Lifestyles (Natsal-3), a population-based survey conducted in 2010-2012 [22,23]. 15.4% individuals reported zero partners, and amongst the remainder the distribution of number of partners approximately followed a power-law distribution (Fig 1A). We used Bayesian inference to estimate the exponent γ of this power-law distribution, and found a posterior mean of γ =1.81 (95% credible interval: [1.69,1.96]). This is comparable to estimates calculated based on the previous Natsal data, collected in 2000 and 1990, and the London Gay Men’s Sexual Health Survey [24], which were 1.57 (95% CI: [1.43, 1.72]), 1.75 (95% CI: [1.57, 1.95]) and 1.87 (95% CI: [1.80, 1.94]) respectively [14].

**Fig 1.**
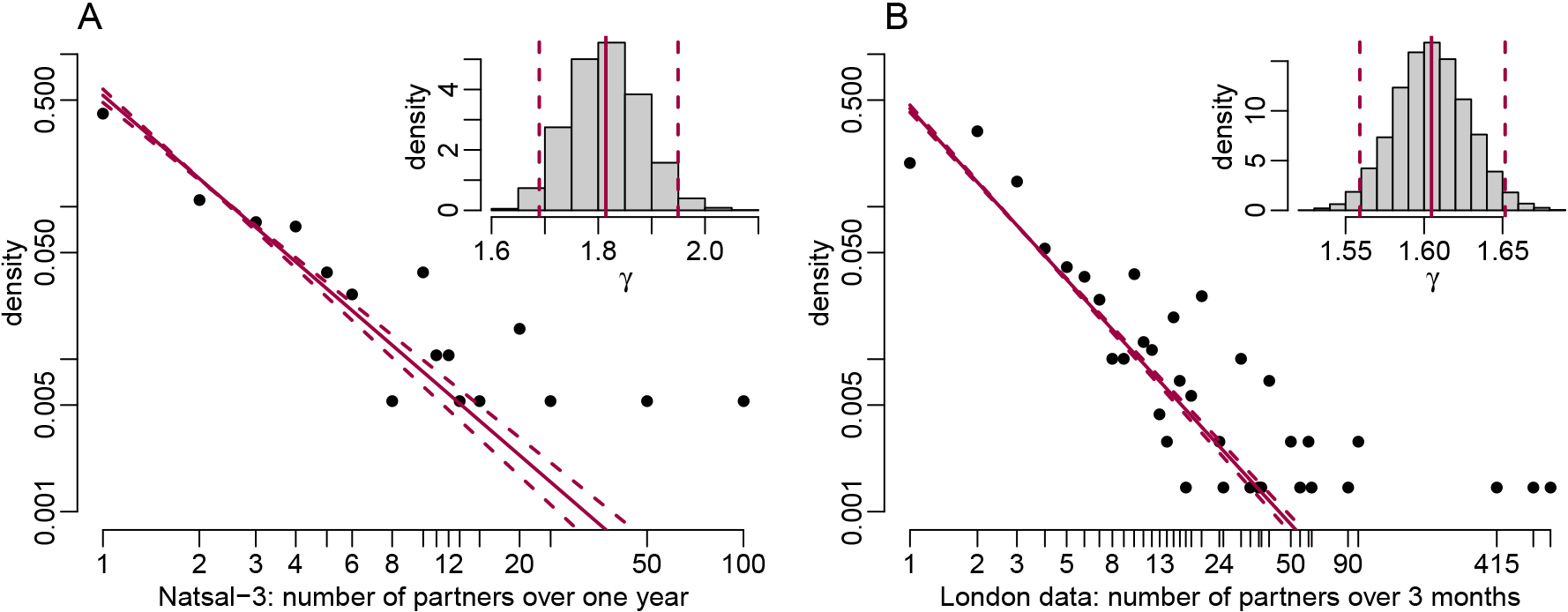
Double logarithmic plot of cumulative degree distribution of number of partners for UK MSM **A:** reported over 1 year by Natsal-3 respondents. **B:** reported over 3 months in GRASP data from London. Inset histograms show posterior distributions of γ for each dataset with mean and 95% credible intervals in red

We performed the same analysis based on data collected in 2004 as part of the national Gonococcal Resistance to Antimicrobials Surveillance Programme (GRASP) run by Public Health England (PHE) from individuals diagnosed with gonorrhoea in London [25,26]. Fewer individuals had only one partner in the last three months than would be expected under a power-law distribution, however a power-law tail was observed for MSM having more than one partner. The inferred scale-parameter γ for gonorrhoea infected individuals was significantly lower than that in the Natsal-3 data at 1.60 (95% credible interval: [1.56, 1.65]) with non-overlapping credible intervals (Fig 1B).

### Simulation of dynamic network model

We developed a new algorithm to simulate dynamic sexual networks in which relationships are being formed and broken down over time. To incorporate sexual behaviour heterogeneity, each individual in the network is characterised by a parameter λ that represents the propensity to make and break relationships. This λ parameter is analogous to the fitness property used in a previously published method to generate static power-law networks [17, 18]. The mathematical properties of our method (see Methods section) imply that the number of partnerships in which individuals were involved over one year is power-law distributed.

To demonstrate the ability of our algorithm to simulate realistic networks, we generated dynamic sexual networks of size *N* =10,000 over one year using a power-law exponent γ equal to 1.7, 1.8 and 1.9 (Fig 2, Fig S1). The network size was chosen to represent MSM aged between 15 and 65 in a UK city such as Brighton or central Manchester [27,28]. Our algorithm also requires to set the parameter *k*_0_ which determines the proportion of individuals that do not have a sexual partnership during the year. Using values of *k*_0_ equal to 0.4, 0.5 and 0.6 respectively, we were able to produce networks exhibiting a power-law distribution of partnerships and proportion of individuals with zero partners over one year that were comparable to the 15.4% proportion in the Natsal-3 data (Fig 2). Finally, a third parameter *ϕ* in our method determines the rate of partnership breakdown, which in turn decreases the level of partnership concurrency in the network without affecting the distributions of partner numbers.

**Fig 2.**
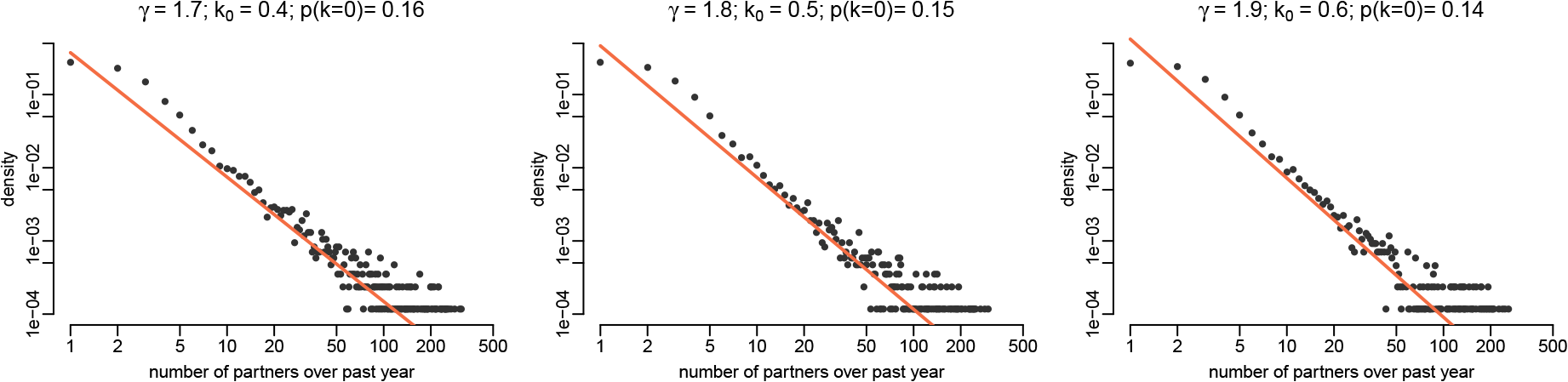
Double logarithmic plot of degree distributions of partners in last year generated with differing parameters γ and *k*_0_. The orange lines represent the desired degree distribution *p*(*k*) = *ck*^−*γ*^. The proportion of individuals having no partners in the last year is shown in the title of each plot as *p*(*k* = 0).

### Effect of network model on simulated gonorrhoea outbreaks

To assess the importance the underlying sexual network structure and dynamism in the way gonorrhoea outbreaks spread, we performed a comparison of simulated gonorrhoea outbreaks on three types of networks (Fig 3A): a fully connected network, a static power-law network and our new dynamic power-law network. On each type of network we used the same model of gonorrhoea outbreak, adapted from a recent study [29] as illustrated in Fig 3B and described in the Methods section.

The flow parameters were calibrated for each of the three types of network in order to produce for each type of network outbreaks of the same realistic size over a year (cf Methods section). The resulting parameter values are summarised in Table 1 with no significant difference between the three models for any parameter except the rate of transmission per partnership *β*, which takes widely different values as expected. From the resulting simulations we analysed the offspring distribution, defined as the number of onward transmissions attributable to each case infected in the first year (Fig 4A) and we extracted the generation times, defined as the length of time from initial infection to onward transmission (Fig 4B). Both the offspring distribution and the generation time distribution exhibited important differences depending on the underlying type of sexual network considered (Fig 4).

The basic reproduction number is defined as the mean of the offspring distribution in a fully uninfected population. For all three network structures (fully connected, static, and dynamic) the basic reproduction numbers were around 1.2 with overlapping 95% ranges (Fig 4A, X axis). This equality is due to the calibration of the models, which required outbreaks to be of similar sizes. However, we found that the variance in the simulated offspring distributions differed depending on network structure (Fig 4A, Y axis). Simulated outbreaks in both static and dynamic power-law networks had greater variance in the offspring distribution than the fully connected networks (4.2; 95% range: [1.9, 7.7]), due to the effect of heterogeneity in contact patterns. However, dynamic partnership formation and dissolution partially mitigated the impact of the network structure on the offspring distribution. The sample variance of the offspring distributions in the static power-law network (10.9; 95% range: [5.7, 20.3]) was on average greater than in the dynamic network (6.8; 95% range: [3.4, 12.0]), suggesting that adopting a static power-law network in disease models would overstate the importance of super-spreading events.

**Fig 3.**
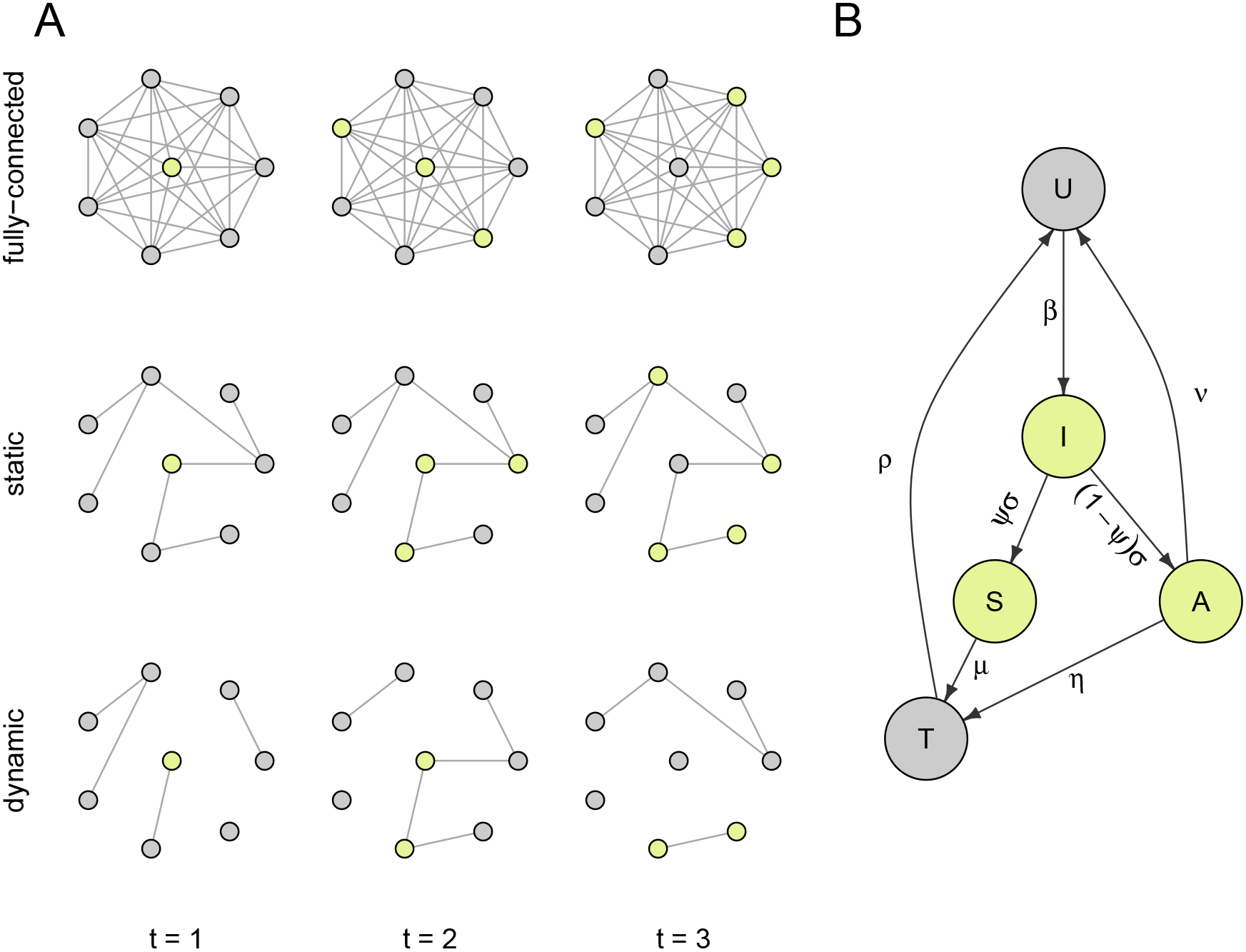
**A :** Illustration of gonorrhoea transmission over time through the three network structures: fully connected, static and dynamic. **B:** Flow diagram of transmission model with rates of transition between infection states. Uninfected individuals (*U*) become infected after sexual contact with contagious individuals. Infections initially pass through an incubation period (*I*), before either developing symptoms (*S*) or remaining asymptomatic (*A*). Symptomatic individuals seek treatment, and asymptomatic infections identified by screening are also treated (*T*).

**Table 1.** Parameter notations, input range (cf Appendix S3) and range of parameters that resulted in outbreaks persisting at least one year in at least 20% of simulations with at most 400 total diagnoses per year on average for each network structure

|  | Parameter description | Unit | Input range | Source |
| --- | --- | --- | --- | --- |
| $\beta$ | rate of transmission within each partnership | year<sup>-1</sup> | [0, 60] | fitting |
| $\psi$ | infections that become symptomatic | % | [40, 95] | [30–32] |
| $1/\sigma$ | duration incubation period | days | [2, 10] | [33–37] |
| $1/\nu$ | duration of carriage | days | [60, 550] | [38–42] |
| $\eta$ | rate of screening when asymptomatic | year<sup>-1</sup> | [0.5, 4] | [43–45] |
| $1/\mu$ | time to symptomatics seeking treatment | days | [1, 30] | [46] |
| $1/\rho$ | time to recovery following treatment | days | [5, 10] | [47] |

|  | accepted parameter mean [95% range] |  |  |
| --- | --- | --- | --- |
|  | fully connected | static | dynamic |
| $\beta$ | 1.8 [1.1, 3.9] $\times 10^{-3}$ | 0.11 [0.08, 0.15] | 24.0 [11.0, 53.9] |
| $\psi$ | 55 [40, 83] | 63 [41, 88] | 71 [41, 94] |
| $1/\sigma$ | 4.8 [3.0, 8.0] | 4.7 [3.0, 7.8] | 4.6 [2.9, 7.8] |
| $1/\nu$ | 223 [113, 459] | 167 [93, 392] | 166 [96, 385] |
| $\eta$ | 1.6 [0.8, 2.8] | 2.2 [1.0, 3.7] | 2.1 [1.0, 3.6] |
| $1/\mu$ | 2.8 [1.2, 21.1] | 2.7 [1.1, 15.4] | 2.7 [1.1, 13.2] |
| $1/\rho$ | 6.9 [5.7, 8.5] | 6.9 [5.7, 8.4] | 7.0 [5.8, 8.5] |

The distribution of mean generation times in the simulations is also affected by the underlying network structure (Fig 4B, X axis). Outbreaks in the dynamic network structure have a mean generation time of 63 days (95% range: [33, 101]). Generation times are overestimated when partnership dynamics are ignored, as in the case of the static power-law network structure (77 days; 95% range: [49, 124]), an effect which is exacerbated when heterogeneity in sexual activity levels is omitted, as in the fully connected network (109 days; 95% range: [54, 168]). In order to maintain persistence of the outbreak at realistically low prevalence, as is observed in gonorrhoea, outbreaks in the fully connected network overstate the proportion of asymptomatic infections compared to the dynamic network, 44.6% (95% range: [17.2%, 59.7%]) compared to 29.5% (95% range: [6.4%, 58.8%]) [5]. This larger untreated asymptomatic reservoir also serves to increase the variance of the simulated generation times from 6,960 (95% range: [2,100, 17,530]) in the dynamic networks to 13,500 (95% range: [3,850, 26,200]) in the fully connected networks (Fig 4B, Y axis).

**Fig 4.**
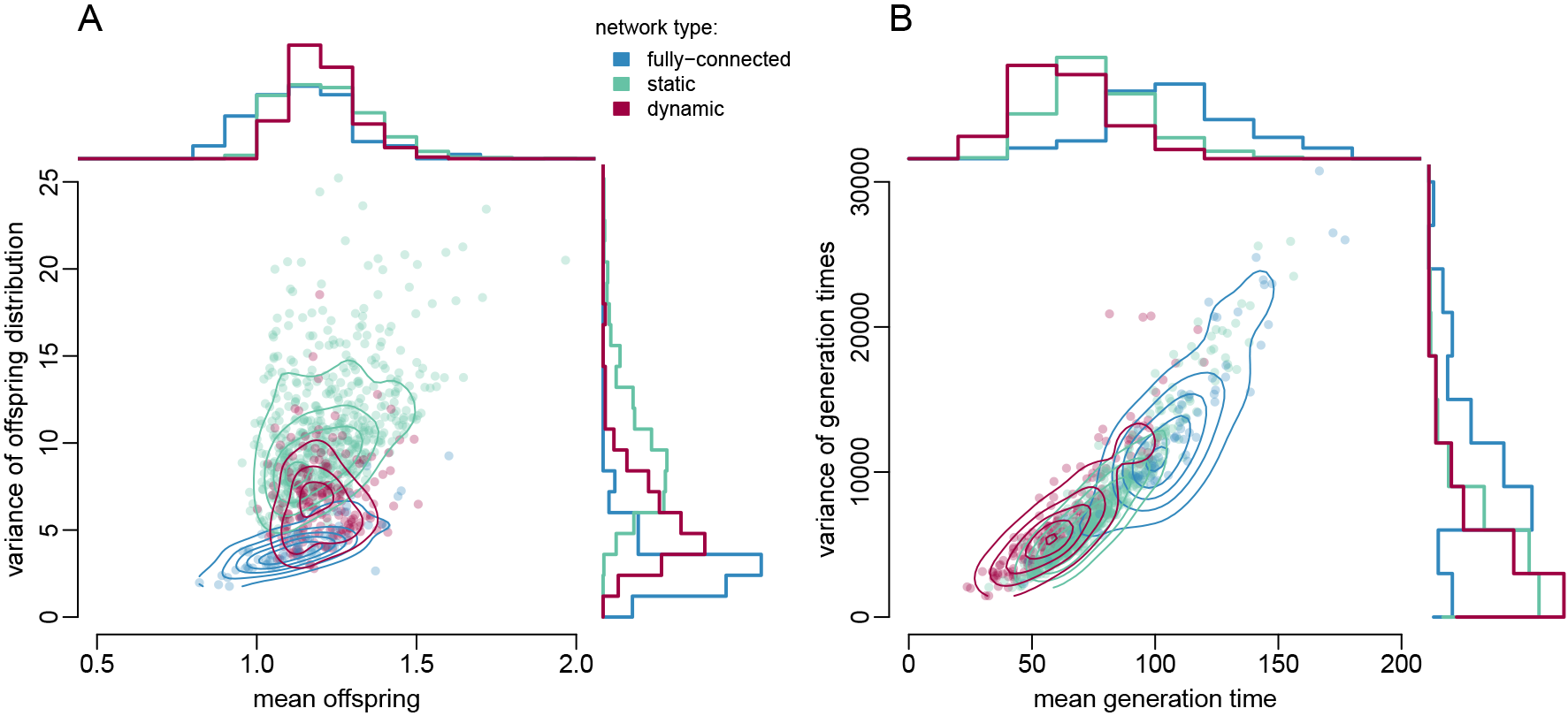
Scatter plots with overlaid density contours and marginal histograms for the mean and variance of **A:** the offspring distributions, and **B:** the distribution of generation times, for infections contracted in the first year of the outbreak. Simulations under the fully connected, static, and dynamic network structures are shown in blue, green and red respectively.

For both the static and dynamic power-law networks we investigated the relationship between the number of sexual partners that an individual has over one year, the probability of becoming infected, and the number of transmission events arising from those individuals who become infected. Fig 5 shows the proportion of infected individuals, the probability of an individual becoming infected in first year and the mean onward transmissions for infected individuals, split by the number of partners over one year. The proportion of infected individuals having fewer than three partners per year was lower than would be expected under a power-law distribution for both static and dynamic networks, however the distribution exhibited power-law behaviour for more highly active infectees (Fig 5A). This is similar to the pattern exhibited in the GRASP London data (Fig 1B). Compared to the dynamic network the static network structure appears to overestimate the burden of infection in individuals with more than 11 partners per year, while underestimating the burden in individuals with fewer partners.

Under the static network structure an individual’s probability of infection increases linearly with their annual number of partners (Fig 5B). When we allow for partnership dynamics, the probability of infection increases linearly at first, albeit at a slower rate than the static network, then levels off in individuals having more than five partners per year, eventually approximating the probability of infection observed in a fully connected network. This suggests that a static network structure may underestimate the risk of infection for individuals with few sexual partners.

**Fig 5.**
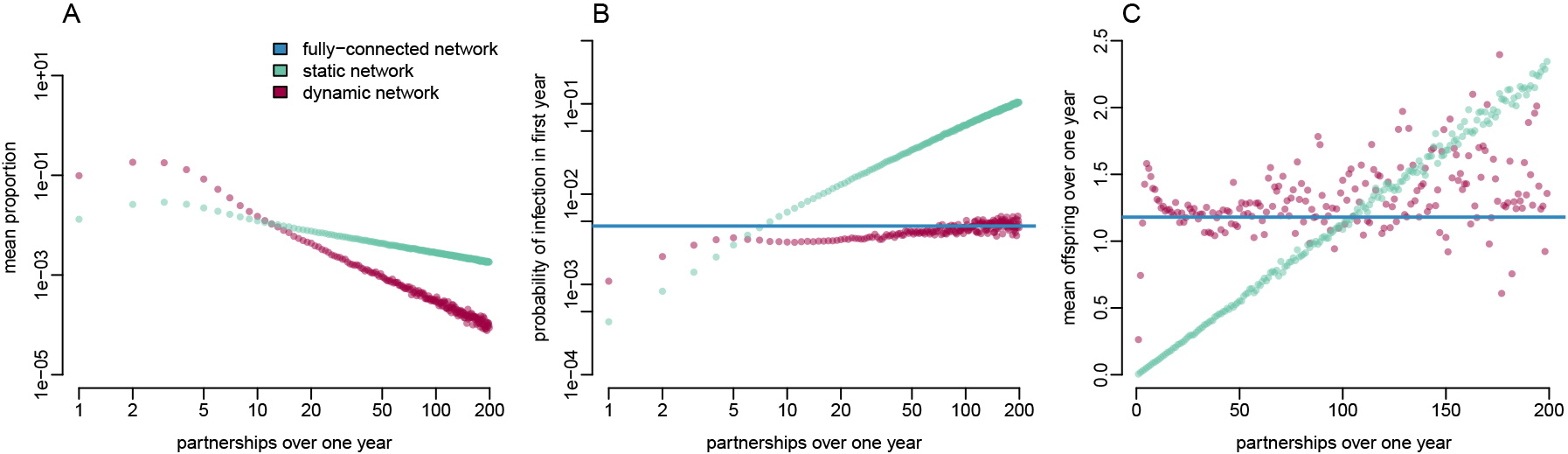
**A:** Proportion of infected individuals split by number of partners over one year **B:** Probability of an individual becoming infected in first year given their total number of partnerships. **C:** Basic reproduction number - mean onward transmissions for infected individuals with a given number of partnerships.

There is a similar relationship between the expected number of onward transmission events from individuals and their number of partners over the year in the static and dynamic network structures (Fig 5C). An infected individual’s expected number of transmissions in the static network increases linearly with their total number of partnerships. For individuals with up to five partners per year, the dynamic network also shows a strong linear relationship between the expected offspring and number of partners. However, for individuals with more than five partners per year the relationship is less strong with a much greater variance in mean number of offspring. In the dynamic network the expected number of offspring is greater than one in individuals with at least three partners, whereas the mean offspring per person in the static network only becomes greater than one in individuals having more than 90 partners. The static network therefore likely overestimates the importance of very highly active individuals in maintaining transmission.

### Impact of interventions

Using the dynamic network model we investigated the impact of a range of interventions and preventative measures against gonorrhoea, including: 20% increased condom use (resulting in a reduction in the transmission rate *β*), 20% increased sexual health screening (an increase in parameter *η*), and the impact of a hypothetical gonorrhoea vaccine, deployed either at random to 20% of the network or targeted to the 20% of individuals with the highest propensity to form partnerships (λ). Recent estimates suggest that the meningococcal B vaccine may be 31% [21%, 39%] effective against gonorrhoea [48]. The impact of vaccinating 20% of individuals at random with a vaccine that is 100% effective is comparable to vaccinating 65% [51%, 95%] of individuals with a vaccine of similar effectiveness. We assessed the one-year impact of these four measures on the probability of a outbreak stemming from a single introduction of gonorrhoea into a dynamic sexual network (Fig 6A), the total number of gonorrhoea diagnoses (Fig 6B), and the number of sexual health clinic visits from both screening and symptomatic treatment-seeking (Fig 6C).

**Fig 6.**
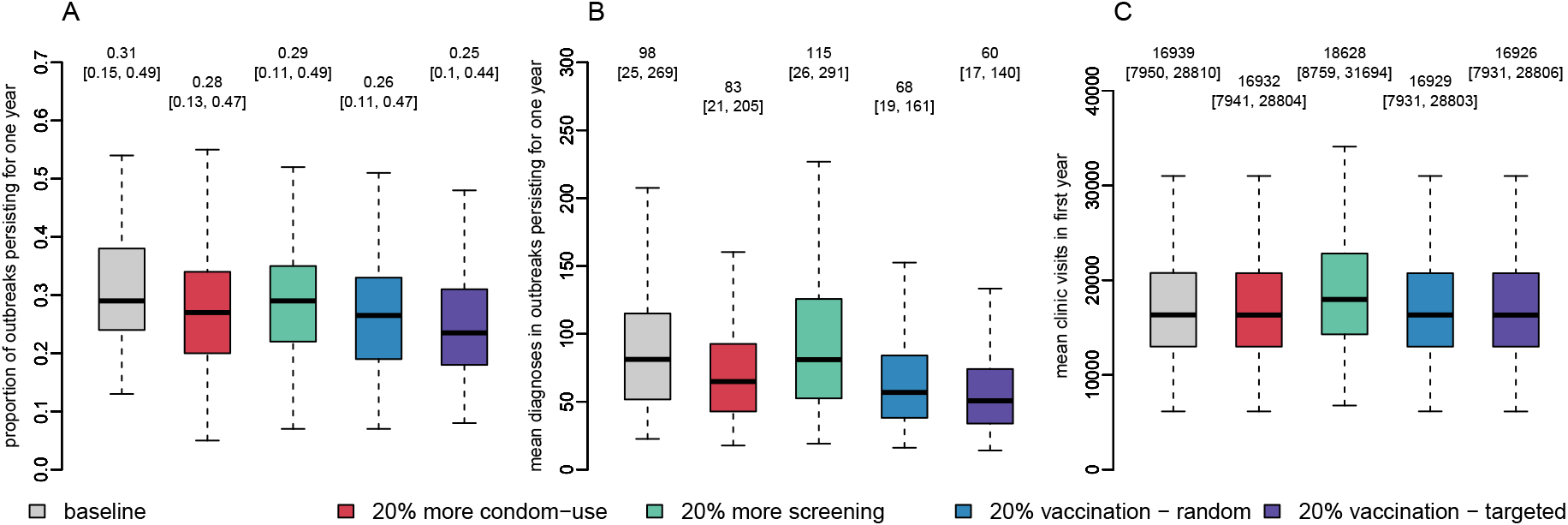
**A:** Proportion of outbreaks persisting for at least one year. **B:** Mean number of gonorrhoea diagnoses in first year. **C:** Mean number of clinic visits in first year.

The baseline proportion of simulated outbreaks persisting for at least one year from a single introduction of gonorrhoea was 31% (95% range: [15%, 49%]). All of the interventions we considered reduced the probability of an outbreak, with vaccination having the greatest impact; a fully effective vaccine administered to 20% of individuals in a randomised strategy reduced the probability of an outbreak by around a third to 21% (95% range: [7%, 40%]), a targeted strategy had a greater effect, reducing the probability to 19% (95% range: [5%, 36%]). The non-vaccine interventions had less of an impact. Increasing condom usage by 20% reduced the probability of an outbreak by around 10% to 28% (95% range: [13%, 47%]); A 20% increase in the rate of screening for asymptomatic cases had a similar effect, reducing the probability of an outbreak to 29% (95% range: [11%, 49%]).

The expected size of the visible outbreak in a population of 10,000, as measured by the total number of infected individuals diagnosed and receiving treatment, was reduced by a fifth from 98 (95% range: [25, 269]) to 79 (95% range: [19, 217]) with a 20% increase in condom-use, and could be halved using vaccination: down to 47 cases (95% range: [14, 114]) using the randomised strategy and 45 cases (95% range: [14, 105]) by targeting the most sexually-active individuals. However, a 20% increase in the screening rate resulted in a 5% increase in the visible outbreak size to 103 cases (95% range: [25, 295]), due to more asymptomatic cases receiving treatment.

There was a similar pattern in the burden of sexual health services, while increased condom use decreased the total number of clinic visits by 11% from 16,939 (95% range: [7,950, 28,810]) to 15,053 (95% range: [7,064, 25,602]). A 20% increase in the rate of screening, both of uninfected and asymptomatically infected individuals increased the total clinic visits by 20%, because the majority of testing is prompted by screening rather than symptomatic treatment seeking. The number of sexual health clinic visits remained stable in the vaccine scenarios, however the financial and administrative cost of initiating either a targeted or randomised vaccination programme must be considered once a vaccine candidate has been developed. It is important to note that while the targeted strategy is more effective, requires the ability to identify and vaccinate the 20% most sexually active individuals in a given population. It is unclear how effectively this could be done in practice, but could perhaps be offered at the GUM clinic at the same time as testing.

## Discussion

Real-world sexual networks are dynamic by nature, and we have developed a method that can reproduce observed power-law distributions of numbers of sexual partners in the last year in a dynamic network, which accounts for heterogeneity of individual behaviour in the way relationships form and break down. Our model allows the user to specify the power-law distribution via the exponent γ, to vary the proportion of individuals having no partners via the parameter *k*_0_, and to set the average length of partnerships, via the parameter *ϕ*. Varying the length of partnerships for a given degree distribution impacts the pattern of partnership concurrency in the network. The longer the average partnership, the higher the degree of concurrency. We implemented this dynamic simulation algorithm into a R package called simdynet which is freely available at https://github.com/lwhittles/simdynet.

Taking an outbreak of gonorrhoea as a case study, we found that failing to allow for sexual network structure (i.e. using the fully-connected network)resulted in an overestimation of the duration of carriage and asymptomatic reservoir. When network structure, but not dynamics of sexual partnership formation and breakage, was accounted for (i.e. using the static network) the model overstated the likelihood of super-spreading events and the burden of disease among individuals with high numbers of partners, compared to a dynamic model. While it is important to take heterogeneity into account, the traditional formulation of a core group [5, 20] might approximate the true transmission dynamics of gonorrhoea better than using a static power-law network. Our findings add support to previous modelling work that suggested that having more sexual partners does not greatly impact the rate at which antibiotic resistant gonorrhoea can spread [8].

We used our realistic dynamic power-law network model to investigate the impact of a range of interventions against gonorrhoea, including increased condom use, more frequent screening and immunisation. Our results confirm that vaccination shows great potential to reduce the burden of gonorrhoea [49]: if a random 20% of individuals were immune, then the probability of outbreaks persisting at least a year would be reduced by 16% with the outbreak size reduced on average by 31%. (Fig 6). Such a level of protection could be achieved either through vaccination of a small portion of the population with a highly effective vaccine, or by widespread use a less effective vaccine. For example, a recent retrospective case-control study has shown that the MeNZB vaccine against meningitis is cross-protective against gonorrhoea, with an estimated effectiveness between 20% and 40% [50, 51].

The dynamic sexual network we implemented is highly realistic, but also too computationally expensive to be used in many applications. For example if the population under study is very large, keeping track of every partnership formation and break down is clearly inefficient, especially for the study of the early stages of an outbreak of a new resistant strain in which only a small subset of individuals are being affected. However, we have computed from the full dynamic model the offspring distribution and distribution of generation time (Fig 4). These estimates allow for our model to be approximated as a stochastic branching process [52,53], where infected individuals transmit to a number of secondary cases drawn from the offspring distribution, and the intervals of time between each primary and secondary cases are drawn from the generation time distribution. The advantage of such a model formulation is that it is much simpler than the full dynamic sexual network we described in this paper, but retains the same basic properties in terms of the transmission process. Furthermore, a branching model is at the basis of several recently developed methods to reconstruct transmission trees from genomic data, such as `outbreaker` [54], `TransPhylo` [55,56] and `phybreak` [57]. Our estimates of the generation time distribution and offspring distribution therefore pave the way for these genomic epidemiology methods to be applied to the reconstruction of transmission in gonorrhoea outbreaks [38,58–60].

We estimated that on average the mean and variance of the generation time distribution were equal to 63 days and 6980 days^2^, respectively (Fig 4B), which can be emulated using a Gamma distribution with shape and scale parameters equal to 0.57 and 110.48, respectively. The resulting 95% quantile range of the generation time stretches up to 298 days, which is in good agreement with an analysis of genomes from pairs of known sexual contact, in which the greatest observed time to most recent common ancestor was 8 months [38]. Since gonorrhoea can often remain asymptomatic, any outbreak reconstruction would need to account for the possibility of unsampled cases acting as intermediates in the transmission chains [38,54,56]. Accurate inference of unsampled cases requires in turn a good prior knowledge of the generation time distribution like the one we estimated here based on a dynamic power-law network. Both the mean and variance of the generation time distribution are overestimated when considering a fully-connected or static network (Fig 4B) which would likely result in an underestimation of the role played by unsampled cases.

The offspring distribution was estimated to have a mean and variance equal to 1.2 and 6.8, respectively (Fig 4A). The mean corresponds to the basic reproduction number and its value results from the conditions we imposed on the frequency and size of outbreaks. Since the variance is greater than the mean, the offspring distribution is over-dispersed compared to a Poisson process, which indicates the presence of super-spreaders [61–63], although this transmission heterogeneity is not as pronounced as would be implied by an unstructured or static network model (Fig 4A). In a branching model, overdispersion can be implemented using a Negative-Binomial distribution for the number of offspring, which in this context is often parametrised in terms of its mean and dispersion parameter *k*, with lower values of *k* indicating more over-dispersion [56,61,64,65]. Here we estimated that *k* = 0.257, which with an offspring mean number of 1.2 gives a 95% quantile ranging up to 9 secondary cases, compared to only 4 cases for a Poisson distribution with the same mean. These estimates of the generation time distribution and offspring dispersion parameter pave the way for future studies of genomic epidemiology in gonorrhoea outbreaks.

## Materials and methods

### Estimation of power-law exponents from real data

The third National Survey of Sexual Attitudes and lifestyles in the UK (Natsal-3), was conducted between September 2010 and August 2012 in 15,000 adults aged between 16 and 74 [22,23]. We extracted the number of same-sex partners over one year for the 188 men who reported sexual contact with another man within the past five years. 15.4% of MSM reported having no same sex partners over the past year.

In addition to the Natsal data we examined the number of partners reported by 691 MSM diagnosed with gonorrhoea in a collection of 2,045 isolates sampled between June and November 2004 from 13 major sexual health clinics throughout London as part of the national Gonococcal Resistance to Antimicrobials Surveillance Programme (GRASP), run by Public Health England (PHE) [25]. PHE has produced a GRASP report annually since 2000 to monitor trends in resistance and susceptibility to the drugs used to treat gonorrhoea in England and Wales, which is used to inform national treatment guidelines and strategy. The GRASP data from London represent 54% of the 3,754 cases reported across the city at that time [26,38,66].

We fitted the power-law distribution using Bayesian inference, implemented via a Monte Carlo Markov Chain, to obtain obtained posterior estimates of γ based on the Natsal-3 and GRASP London datasets, using an uninformative γ ∼ 𝒰[1,10] prior. Five chains with over-dispersed starting points were run for 100,000 iterations after a 10,000 iterations burn-in period and thinned by a factor of 100. The convergence of the MCMC was assessed by visual inspection of the trace plots, and confirmed to have a Gelman-Rubin criterion of < 1.1 [67,68].

### Simulating a dynamic power-law network using vertex intrinsic fitness

#### Network dynamics

We consider a population of *N* sexually active men who have sex with men (MSM). Each individual *j* = 1,…,*N* has a propensity λ_*j*_ to form new partnerships which is randomly drawn from a probability distribution with density *f*(λ), and cumulative density function 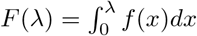. The events that can occur with regard to the network composition are: a new partnership forming (event *a*) or a partnership breakup (event *d*). These events occur as Poisson processes: the (time invariant) rate of individual *j* forming a partnership with individual *l* (given they are not already in a partnership together) is given by the function *a*(λ_*j*_, λ_*l*_). The rate of individuals *j* and *l* breaking up, given they are in a partnership, is denoted *d*(λ_*j*_, λ_*l*_).

The state of each possible partnership is independent from all others, with no limit on the number of concurrent partners. Each possible partnership {*j*, *l*} can be thought of as being ‘on’ or ‘off’ (i.e. is in existence at a particular point in time or not). We can then derive a stationary distribution of the network, and use this to derive the probability at time *t* that each possible partnership exists. By considering the average time spent in each state (on or off) we find that the probability of pair of individuals {*j*, *l*} being in a partnership at time *t* is:

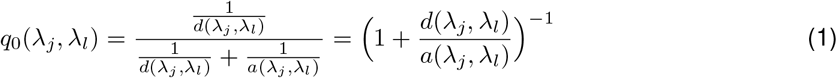

Conversely the probability they are not in a partnership at *t* is simply:

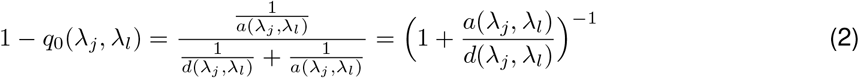

Given that two individuals are not in a partnership at time *t* the time to forming a partnership, is distributed Exp(*a*(λ_*j*_, λ_*l*_)), so the probability of them forming a partnership in the next year is 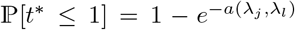. Combining these two cases, we can calculate the probability that two individuals have been in a partnership over the last year:

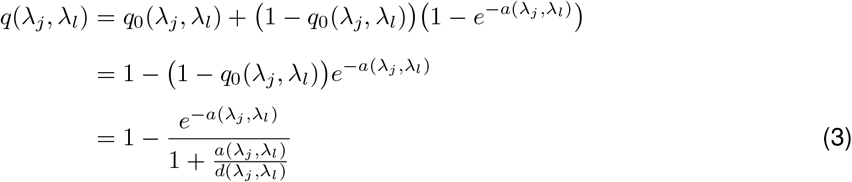

#### Analytical expression of the degree distribution

Let *k*_*j*_ denote the degree distribution of individual *j*, that is the number of partnerships involving *j* that have been active at some point over the past year. The probability that individual *j* was in partnership at some point with an individual *l* of unknown λ_*l*_ is equal to:

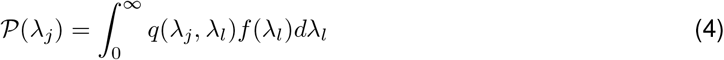

The distribution of *k*_*j*_ given λ_*j*_ is therefore 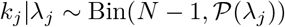, and its expectation is:

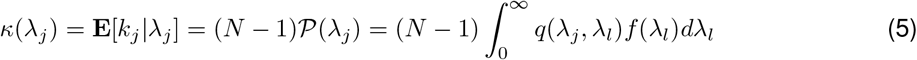

Since *κ* (.) is a continuous, monotonically increasing function of λ, we can use the method of transformations to find the degree distribution *p*(κ):

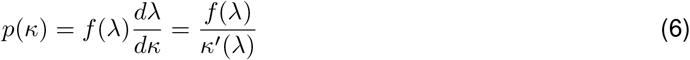

#### Power-law distribution of partners over a year

We require the network of partnerships over the last year to have a power-law degree distribution, so:

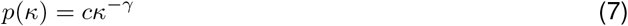

The constant *c* is determined by the fact that 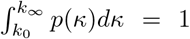, where *k*_0_ = lim_λ→0_*κ*(λ) and *k*_∞_ = lim_λ→∞_ *κ*(λ). For γ ≠ 1 (in accordance with observations from real-world networks) we have:

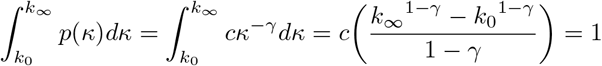

So

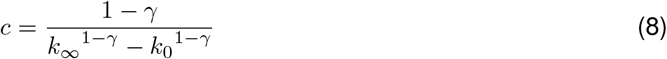

While *k*_0_ is an input of the model, *k*_∞_ is chosen to be as large as possible, subject to the condition that lim_λ_*j*_,λ_*l*_→∞_ *q*(λ_*j*_, λ_*l*_) ≤ 1. We equate Eqs 6 and 7 and rearrange to obtain an expression for the density of λ in terms of the degree distribution:

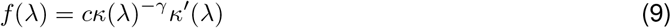

Integrating both sides between 0 and λ we obtain:

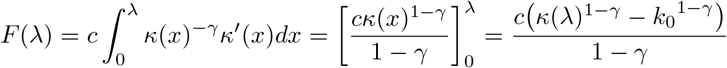

And after rearrangement:

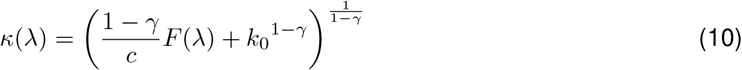

Following [18], we consider the special case where:

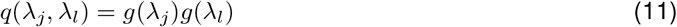

Substituting Eq 11 into 5 we obtain:

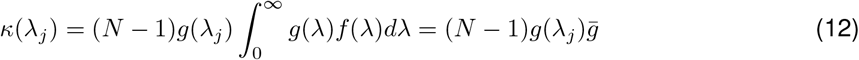

where *g̅* = **E**[*g*(λ)]

Equating Eqs 10 and 12, then multiplying by *f*(λ) and integrating between 0 and ∞ we obtain:

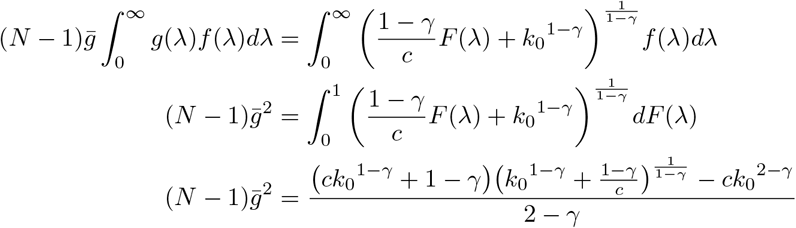

So

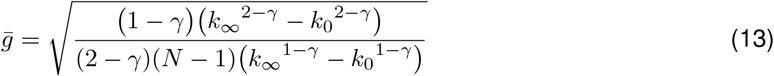

Rearranging 12 we obtain:

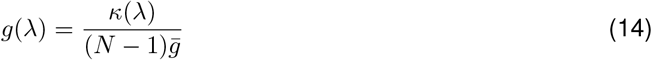

Substituting Eqs 10 and 13 into 14 we obtain:

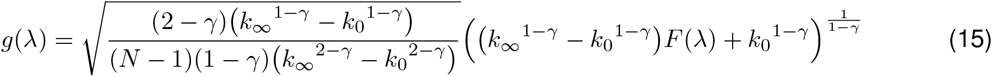

Combining Eqs 15 and 11 we deduce the function *q*(λ_*j*_, λ_*l*_) required to the network to follow a power-law distribution. The functions governing partnership formation and dissolution can then be inferred by considering the simple case where *d*(λ_*j*_, λ_*l*_) = *ϕa*(λ_*j*_, λ_*l*_) where *ϕ* is a constant so Eq 3 becomes

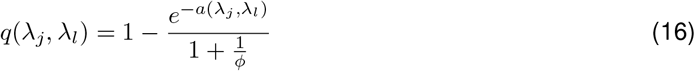

We set the minimum rate of partnership breakup 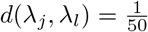, so that the longest partnerships in the network are Exponentially distributed with a mean of 50 years. Applying this constraint and rearranging Eq. 16 we obtain:

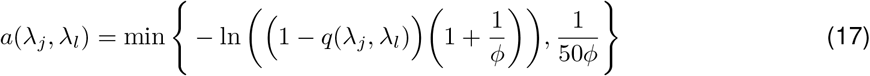

We deduce an algorithm for the simulation of a static snapshot of the network over a year (Appendix S1) and an algorithm for the simulation of the dynamic network (Appendix S2). Both algorithms were implemented in a R package called `simdynet` which is freely available at https://github.com/lwhittles/simdynet.

### Simulation of gonorrhoea outbreaks

In order to investigate the patterns of transmission under different network structures, we consider a stochastic individual-based model of gonorrhoea, adapted from [29]. Individuals are initially uninfected (*U*) and become infected probabilistically at rate *β* due to sexual contact with an contagious individual as dictated by the underlying sexual network structure. Infected individuals initially pass through an incubation period (*I*) which they leave at rate *σ*. A proportion *ψ* of those infected then develop symptoms (*S*), whereas the remainder enters an asymptomatic stage (*A*). In men, gonococcal infection can occur in the rectum, pharynx and/or urethra, resulting in different rates of onward transmission and probabilities of developing symptoms [69]. We do not explicitly model separate anatomical sites of infection, therefore the rate of transmission, *β*, and the likelihood of developing symptoms, *ψ*, should be seen as an average for any infection site. Asymptomatic individuals (*A*) undergo screening and receive treatment at rate *η*, otherwise recovery from asymptomatic infection happens (either naturally or following unrelated antibiotic treatment) at rate *ν*. The symptomatic individuals (*S*) seek treatment at rate *μ*. Individuals who have been treated recover from the infection and become uninfected again at rate *ρ*. The contagious population is denoted *C* = *I* + *S* + *A*, since individuals in treatment are assumed either to no longer be contagious or to abstain from sexual activity in accordance with treatment guidelines [47].

### Model calibration

Using the Gillespie algorithm described in Appendix S2 we generated dynamic sexual networks exhibiting a power-law distribution of partnerships and proportion of individuals with zero partners over one year that were comparable to the Natsal-3 data. A thousand sets of parameters were sampled by Latin Hypercube Sampling from input ranges selected based on published sources (cf Table 1 and Appendix S3). For each underlying network structure (fully connected, static and dynamic) we seeded infection 100 times and simulated over three years for each parameter set. Parameter sets that produced outbreaks persisting at least one year in fewer than 20% of simulations were discarded, as were parameter sets that resulted in total diagnoses exceeding 400 cases per year on average. Thus only parameter sets that resulted in realistic gonorrhoea outbreaks were retained.

We considered a suite of interventions and preventative measures against gonorrhoea, including: increased condom use (resulting in a reduction in the transmission rate *β*), increased sexual health screening (an increase in *η*), and the impact of a hypothetical gonorrhoea vaccine, deployed either at random or targeted to the individuals with the highest propensity to form new partnerships (λ). We assessed the one-year impact on the probability of a outbreak stemming from the introduction of gonorrhoea into a dynamic sexual network, the number of sexual health clinic visits from both screening and symptomatic treatment-seeking, and the total number of gonorrhoea diagnoses.

